# Quantifying the antimicrobial activity of CRISPR-Cas9-accA modified Δ*B. subtilis* mutants against *V. harveyi* and *E. Coli*

**DOI:** 10.1101/2021.08.10.455802

**Authors:** Tatiana Hillman

**Affiliations:** Biotechnology, Independent Research, Los Angeles, California, United States

## Abstract

Probiotics are increasingly popular, currently. Probiotics have been described with the ability to treat many disorders of the gastrointestinal tract (GIT) such as irritable bowel syndrome (IBS) and Crohn’s disease. Types of probiotics include bacterial strains from *Lactobacillus* and *Bifidobacterium*. Probiotics can restore balance to gut microbiota by outcompeting pathogenic bacteria for nutrients and secrete antimicrobials to eliminate these bacterial pathogens. However, the viability of most advertised probiotics lose their potency due to being freeze dried into powders during storage or for consuming. Many probiotics become ineffective and produce lower CFUs while traversing through the gastric acids of the digestive system. For these reasons, this study sought to enhance the antimicrobial response of a highly potent probiotic known as *Bacillus subtilis. B. subtilis* has been used to treat many disorders of the gut and secrete many antimicrobials lethal for pathogenic microbes. *B. subtilis* was genetically modified to express CRISPR-Cas9 nuclease deletion of the *accA* gene (Δ*B.subtilis* mutants), which inhibits expression of an essential *accA* gene a part of the fatty acid synthesis (FAS) metabolic pathway. The CRISPR-Cas9-*accA ΔB.subtilis* mutants were co-cultured with *V. harveyi* and *E. Coli*. Bacterial growth, biofilm formation, antimicrobial activity, and antibiotic resistance were quantified. It was found that *ΔB.subtilis* mutants co-cultured with *V. harveyi* and *E. Coli* lessened bacterial growth, amplified biofilm with *V. harveyi*, reduced biofilm formation of *E. Coli*, the co-cultures with the mutants lacked antimicrobial activity, and increased the antibiotic resistance of *V. harveyi* and *E. Coli*. It can be concluded that there is an immense potential for using genetically engineered probiotic strains to enhance the antimicrobial activity of *B. subtilis*, which can amplify the reduction of pathogenic bacteria. However, the safety and frugality of using *B. subtilis* as a probiotic requires further consideration.

## INTRODUCTION

Humans carry a massive amount of microbes, forming the human-microbiome. The human gut consists of over 100- to 1000 microbial species, which maintain and regulate the internal environment of a host [1]. This human-microbiome superorganism contributes substantially to the health of the host. Research into this incredible symbiotic relationship has drawn much attention and research studies [1]. These human gut microbes work to defend against pathogens and affect brain-gut responses. Currently, because of the vast and growing knowledge of the human gut microbiome and the negative effects of dysbiosis, probiotics can restore balance to the ecosystem of the intestines [Martin]. Commensal bacteria can be used as probiotics; therefore, giving greater access to new types of probiotics known as Next Generation Probiotics (NGPs) or Live Biotherapeutic Products (LBPs). Probiotics may offer new and novel preventive therapies [2]. Probiotics are living microbes that provide many health benefits to a host. However, dead bacteria can also have beneficial probiotic properties. *Bifidobacterium* and lactic acid strains of bacteria that show probiotic properties have been used in many foods and in dietary supplements. Probiotics have been demonstrated to protect against digestive disorders such as diarrhea, atopic dermatitis as eczema, and prevent *Clostridium difficile*-associated inflammatory bowel disorders in adults [1].

However, a study examined the viability of a few commercial probiotics as the probiotics transitioned through the GIT. The results showed that the commercial probiotics was reduced by 106-fold in colony-forming units (CFU) after 5 minutes of incubation time in gastric fluids [3]. As a result, there is concern that many other commercial probiotics are less viable and ineffective during food processing, while being stored, and through the probiotics’ traversing the upper GIT [3]. If, even, the probiotics arrive into the colon, they may not be able to colonize the gut microbiome and may exit from the colon into the feces [3]. *Lactobacillus* or *Bifidobacterium* strains are used in commercial probiotics and are consumed by harsh environmental conditions in many food products and in the gastric fluids of the human gut. *Bacillus* strains, Pediococcus, and some yeasts are more effective and suitable probiotic options [3]

This research study focused on using *B. subtilis* more as a probiotic because of its increased effectiveness as an antimicrobial probiotic. Because *Bacillus* has highly effective antimicrobial behavior in the gastrointestinal tract, it has been promoted as a potential probiotic [4]. *Bacillus subtilis* forms spores, is non-pathogenic, and is Gram-positive. *B. subtilis* is found in soils and in the GIT of a few mammals. *B. subtilis* effectively regulates and contains the balance of GIT microflora in a mammalian host [5]. *B. subtilis* can generate an intense protective type of biofilm by stimulating signaling pathways that organize gene expression, encoding the extracellular matrix (ECM) [5]. Current studies report the increased popularity of using *Bacillus* species, such as *B. subtilis*, as probiotics [5]. *Bacillus* species were proven to effectively protect against respiratory infections and other disorders of the GIT such as irritable bowel syndrome (IBS) [5]. However, the types of probiotic mechanisms *Bacillus* species use are still unclear [5]. It seems that *B. subtilis* maintains an auspicious balance of the gut microbiota by increasing *Lactobacillus* (LAB) cell growth and potency [5]. *B. subtilis* bacterial cells produce many different antimicrobial substances, such as lipopeptides, surfactants, inturins, and penguins. For this reason, the current study aimed to investigate an enhancement of *B. subtilis* probiotic antimicrobial mechanisms via synthetic and genetic re-engineering of wild type *B. subtilis. B. subtilis* cells were transformed with CRISPR-Cas9 plasmid vectors (Figure 1).

**FIGURE 1.**
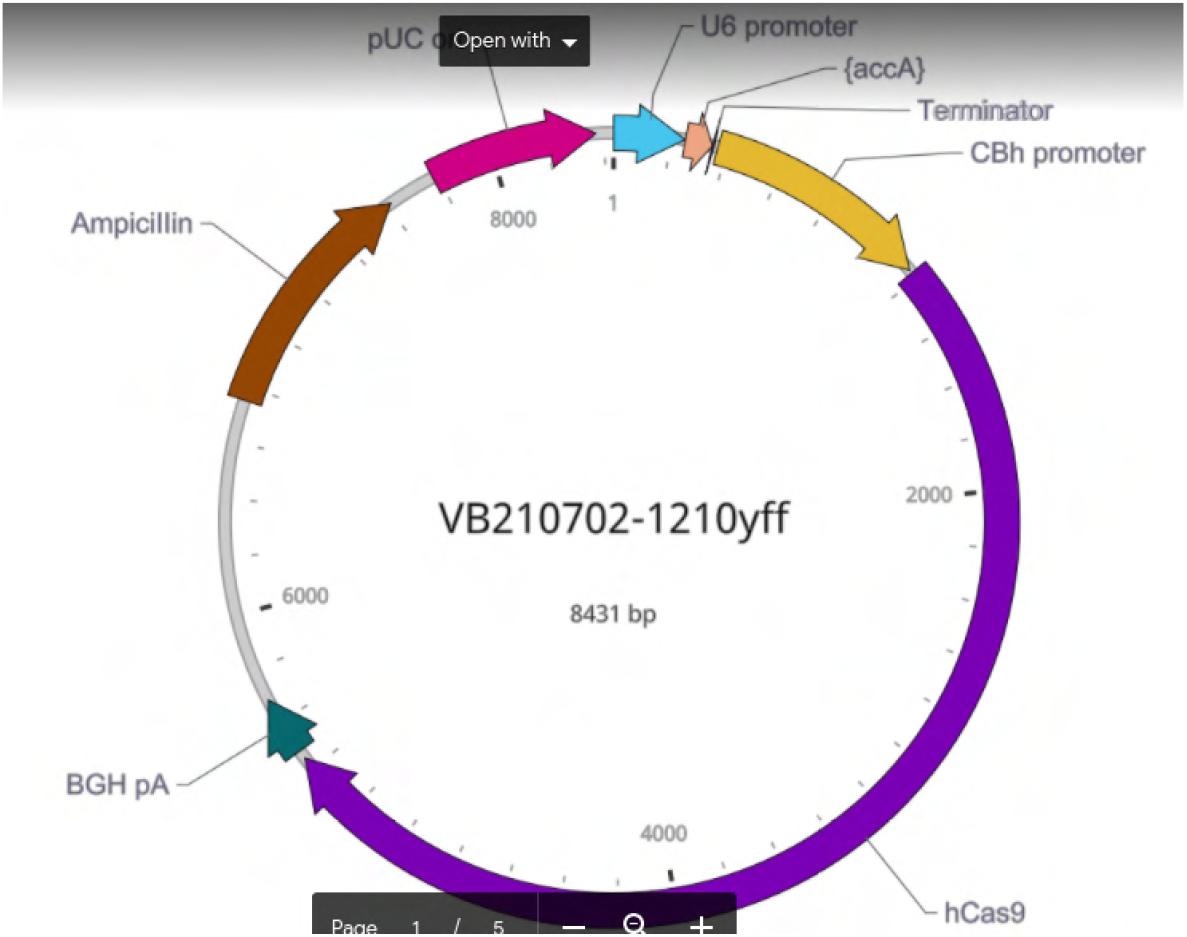
CRISPR-Cas9-accA Plasmid Vector. The vector named pRP[CRISPR]-hCas9-U6>{accA} was designed in VectorBuilder.com. The vector ID is VB210702-1210yff with a total value of 8431 base pairs (bp). The vector was designed to contain a single guide RNA (gRNA) for CRISPR-Cas9 to cleave the accA gene. The gRNA was formed to bind to the sequence of CTGCGTGAAATGTCTCGCCT, which was 20bp of the accA gene. To assemble pRP[CRISPR]-hCas9-U6>{accA} hCas9 was inserted into the vector.

The CRISPR-Cas9 plasmid vectors were designed to express Cas9 nucleases that cleaved and deleted the accA gene. accA codes for the expression of an AccA subunit, a part of the highly conserved and essential acetyl-CoA carboxyl transferase enzyme. Forty-four essential genes are used during synthesis of the cell envelope, which are needed for membrane and cell wall generation. Fatty acids become synthesized into membrane lipids, glycolipids, and into phospholipids. The four genes of accA, accB, accC, and accD combined with acpA and fabD gene products initiate fatty acid synthesis [6]. It has been shown that targeting the ACCase enzyme in bacteria can inhibit its synthesis of fatty acids needed for forming bacterial lipid bilayers and membranes, thereby decreasing the viability and growth of bacteria. The mutant *B. subtilis* was co-cultured with *Vibrio harveyi*, a highly pathogenic bacteria found in many of the fish sources available for human consumption. Through co-culture assays the antimicrobial effects of the CRISPR-Cas9-accA *B. subtilis* mutant on *V. harveyi and E. Coli* were measured and quantified. Overall, the purpose of the present study was to demonstrate the immense potential of using genetically engineered probiotic strains, such as *B. subtilis*, to reduce the presence of pathogenic bacteria and eliminate dysbiosis.

## Methods and Materials

### Bacterial Strains

Bacterial strains included *B. subtilis, V. Harveyi* DSM 6904 (DSMZ), *E. Coli* BL21, and *E. Coli* DH5a. Luria Bertoni broth and agar were inoculated with each strain of bacteria aforementioned. The LB liquid and LB agar media were incubated from 16 hours to 24 hours at the temperatures specific for each bacterium. *B. subtilis* and *V. harveyi* bacterial cells were incubated at 30 degrees Celsius as *E. Coli* cell cultures were grown at 35 to 37°C.

### Plasmid Design and Assembly

The vector named pRP[CRISPR]-hCas9-U6>{accA} was designed in VectorBuilder.com. The vector ID is VB210702-1210yff with a total value of 8431 base pairs (bp). The vector was designed to contain a single guide RNA (gRNA) for CRISPR-Cas9 to cleave the accA gene. The gRNA was formed to bind to the sequence of CTGCGTGAAATGTCTCGCCT, which was 20bp of the accA gene. To assemble pRP[CRISPR]-hCas9-U6>{accA} hCas9 was inserted into the vector. The antibiotic resistant gene for ampicillin was also included in the plasmid assembly. The plasmids were produced at a high plasmid copy number. The fully assembled vectors were assembled and delivered by VectorBuilder. The plasmids were delivered through a glycerol stock solution of *E. Coli* cells. The pRP[CRISPR]-hCas9-U6>{accA} plasmids were purified from the *E. Coli* cells via the BioBasic EZ-10 Spin Column plasmid DNA Kit (BS413-50 preps).

### *B. subtilis* Transformation

The pRP[CRISPR]-hCas9-U6>{accA} plasmids isolated from the *E. Coli* stock of cells were used to transform *B. subtilis* cells, forming ΔB.subtilis mutants. To optimize the *B. subtilis* transformation protocol, *B. subtilis* was cultured for 5 hours unto the exponential phase of growth with an OD600 value of 0.3-0.4. Two millimeters were taken from the culture and centrifuge for 1m at 8,000 rpm, collecting the *B. subtilis* cells. The supernatant was discarded. About 200 μL of LB broth with 2% xylose was added and then vortexed to concentrate the cell suspension at 10x. The 2% xylose was added to the LB. The isolated pRP[CRISPR]-hCas9-U6>{accA} plasmids with ampicillin selective markers were added to the concentrated *B. subtilis* cells with the 2% xylose-LB solution. The *B. subtilis* transformations were incubated for 1 hour at 37° C and then plated. About 20 μL each of the pRP[CRISPR]-hCas9-U6>{accA}*B. subtilis* transformations were inoculated onto two LB agar plates and 160 μL was also plated. The plates were sealed with parafilm and incubated overnight at 37°C. Each LB agar culture plate contained carbenicillin to select for the pRP[CRISPR]-hCas9-U6>{accA} plasmids with the ampicillin selective marker because carbenicillin cannot degrade in high acidity or in excessive heat.

### Co-Culture Assays

The *V. harveyi* and the *E. Coli* bacterial cells were co-cultured with the Δ*B.subtilis* mutants and the wild type (WT) *B. subtilis* to detect any present antimicrobial activity of the Δ*B.subtilis* mutants. After completing the co-cultures, the Δ*B.subtilis* mutant co-cultures with *V. Harveyi* and *E. Coli* BL21/DH5α cells were compared to the co-cultures of the WT *B. subtilis* with the *V. harveyi* and the *E. Coli* BL21/DH5α. To use a co-culture assay and approach, each bacterial strain aforementioned was grown in LB Broth and then plated onto LB agar media. About 5-6 Bacterial colonies from each *E. Coli* BL21/DH5α and *V. harveyi* plate combined with 5-6 colonies from the Δ*B.subtilis* mutant and WT *B. subtilis* plated cultures were inoculated into LB broth. The co-cultures were grown overnight at 30 to 35°C and then analyzed.

### Colony Forming Units (CFU)

To measure the total of colony forming units (CFU), 10μL of each stock of the bacterial strains were pipetted into 1.5 mL Eppendorf centrifuge tubes. About 10μL of the stocks were added to 90μL of LB broth to form the 10^−1^ dilution of each bacterial strain. The dilutions were mixed gently. Then, 10μL of the 10^−1^ dilution was added to 90μL of LB broth to form dilution 10^−2^. The 10μL of 10^−2^ were added to 90μL of LB broth, and then more dilutions were continued in a series to a total of 10^−10^ dilutions. The last dilution and two other dilution series were selected for plating. The plates with the selected dilutions were incubated overnight at 37°C. The number of colonies for each bacterial strain were counted from the cultured agar plates. The number of colonies counted were substituted into an equation for calculating the values of CFU for bacterial cells.

The equations included:

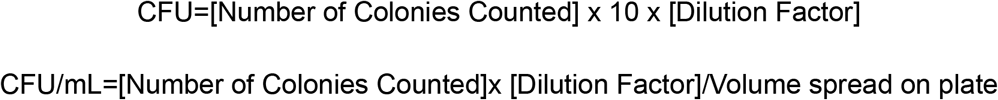

### Kirby-Bauer Disk Diffusion Tests

The co-cultures of *E. Coli* BL21/DH5α and *V. harveyi* with Δ*B.subtilis* mutants and WT *B. subtilis* were then assayed for antibiotic resistance. The Kirby-Bauer Disk Diffusion Tests were used. Carbinicillin, chloramphenicol, tetracycline, and kanamycin were pipetted into sterilized antibiotic disks. Approximately, 5 μL of each of the four antibiotics were pipetted into the antibiotic disks. A lawn of each bacterial strain was inoculated on LB agar plates via the spread plate method. A swab of LB broth liquid cultures from each bacterial strain were spread onto agar culturing media. The antibiotic discs were placed onto the bacterial culture plates. The plates were incubated for 24 hrs at 35°C. The zones of inhibition were measured in millimeters (mm) and then the values interpreted, using a zone diameter interpretive standards chart provided by Sarker et al. [7].

### Biofilm Assay

Five milliliters of LB broth cultures were prepared from each bacterial strain and co-culture. The biofilm of the *E. Coli* and *V. harveyi* strains were used as the controls and were co-cultured with the Δ*B.subtilis* mutants and WT *B. subtilis*. The cultures grew for 20 hours at 37°C. The 1: 100 dilution sets were prepared, totalling to 1mL of the LB/Carb liquid cultures. About 100 μLof the dilution series were pipetted into 4 wells, per bacterial strain, of a 96 well plate. The plate was incubated for 48 hours at 37°C. The plates were shaken to remove any planktonic bacteria. The plates were rinsed with water. All the wells were stained with 125 μL of 0.1% crystal violet solution for 10 min. The 96-well plate was shaken over a tray and the crystal violet was rinsed out into water. The plate was allowed to air dry overnight, and 200 μL of 30% acetic acid was added to all the stained wells to solubilize the crystal violet. The acetic acid was allowed to sit and stabilize for 10 minutes. The mixture of acetic acid and crystal was resuspended by pipetting. The 125 μL of acetic acid, including crystal violets, was transferred from each well into a second 96-well plate.

### Cross-Streak Plate Method

A similar cross-streak method by Lertcanawanichakul and Sawangnop was used [8]. LB agar plates were inoculated with Δ*B.subtilis* mutants and WT *B. subtilis* by applying a single elongated streak of the inoculums to the central most part of each agar plate. The cultures were grown for 2 days of incubation at 37°C. After 2 days, the *E. Coli* BL21/DH5α and *V. harveyi* were streaked on the Δ*B.subtilis* mutant and WT *B. subtilis* plates at a 90° angle or perpendicular to each *Bacillus* species. The cross-streak plates were examined for the size of the inhibition zones measured in mm.

### Statistical Analysis

The p-Values of Colony Forming units for *V. harveyi* samples were analyzed through paired-Two-Tailed t-test. The biofilm assays were analyzed via Two-Way ANOVA. The antibiotic test data results were analyzed by One-Way ANOVA for *E. Coli* samples as paired-t-tests were used to generate p-Values for *V. harveyi* antibiotic resistance sample results. All graphs were constructed through ChartExpo.

## RESULTS

### Bacterial Growth Decreased in V. Harveyi, *E. Coli*, and Δ*B.subtilis* mutant Co-cultures

Co-culture assays were used to detect possible antimicrobial activity between *V. Harveyi, E. Coli*, and Δ*B.subtilis* mutants, which carried pRP[CRISPR]-hCas9-U6>{accA} plasmid vectors. *B.subtilis* was transformed with CRISPR-Cas-9 plasmid vectors targeting the accA gene to inhibit fatty acid synthesis. The goal included observing any enhancement of the Δ*B.subtilis* mutant’s antimicrobial abilities against *V. harveyi* and *E. Coli* target bacterial strains. As a result, after calculating the colony forming units per milliliter (CFU/mL), bacterial growth, in the Δ*B.subtilis* mutants to *V. harveyi* co-cultures, significantly decreased with a paired-t-test Two-Tailed p-Value of 0. *V. harveyi* decreased by an average of 31% after the co-culture assays (Figure 2). The bacterial growth between *E. Coli* and Δ*B.subtilis* mutant co-cultures produced a percent difference of 54%. *E. Coli* yielded 3.02E7 CFU as the co-culture decreased to 1.10E7 CFU. *V. harveyi* had an average CFU of 5.8E11 that then reduced to 3.4E11 CFU during the co-culture assays (Figure 3). Overall, the co-cultures exhibited increased antimicrobial activity. To further detect the antimicrobial behavior of the Δ*B.subtilis* mutants in *V. harveyi* and *E. Coli* co-cultures, the biofilm of the co-cultures were assayed.

**FIGURE 2.**
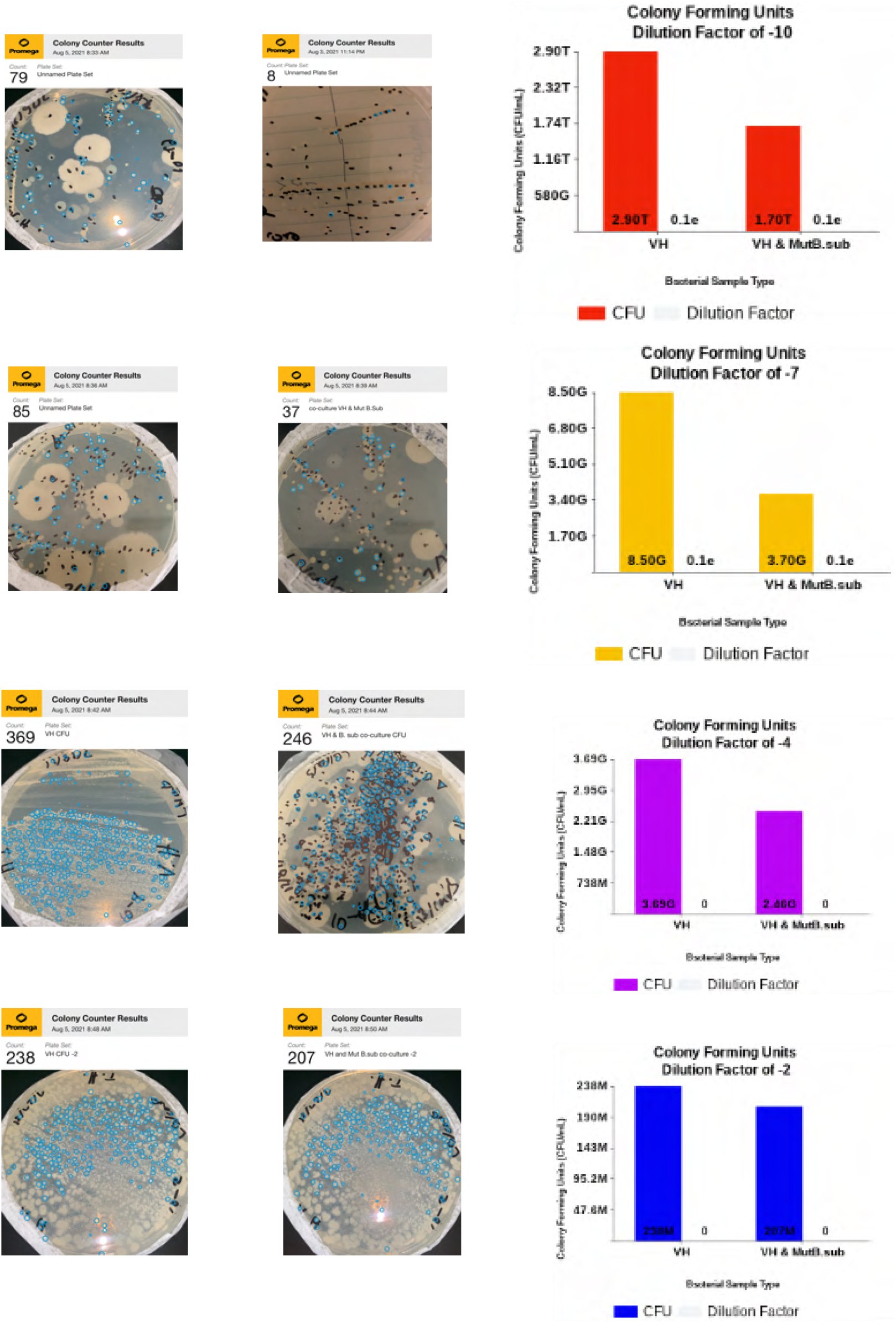
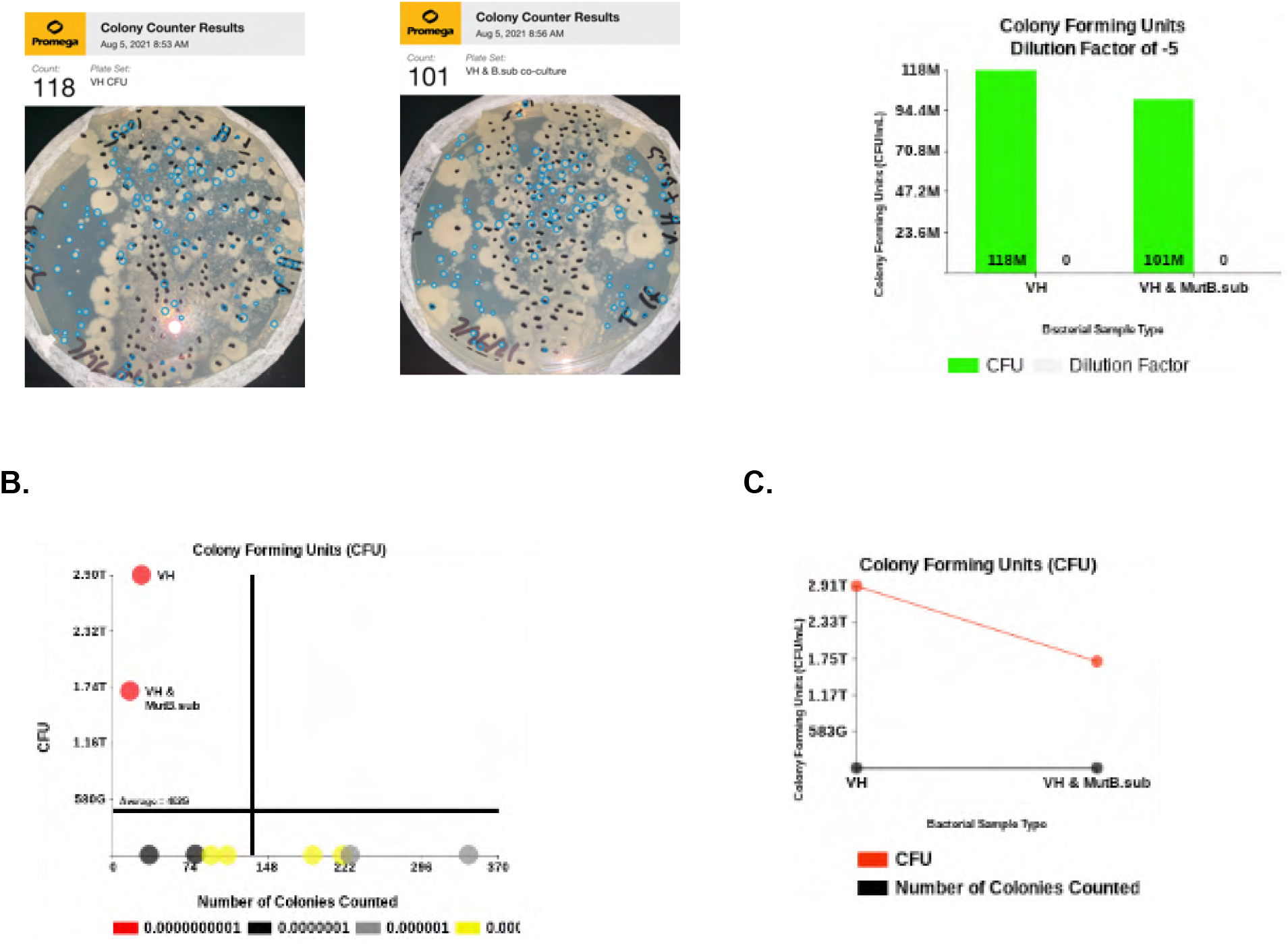
The Colony Forming Units (CFU/mL) After dilution series for each sample were generated, the CFU/mL was allocated. **A.** The dilution series included 10^−10^, 10^−7^, 10^−4^, 10^−2^, and 10^−5^. Each dilution series showed a large decrease in CFU/mL for Δ*B.subtilis* mutant-V.H compared against the *V. harveyi* only bacterial sample culture. **B.** V.H CFUs dwindled from 290 trillion to 1.74 trillion when *V. harveyi* was co-culture with Δ*B.subtilis* mutants. **C.** A decreasing trendline was formed, exhibiting a significant decrease in CFU formed during Δ*B.subtilis* mutant-V.H co-culture assays.

**FIGURE 3.**
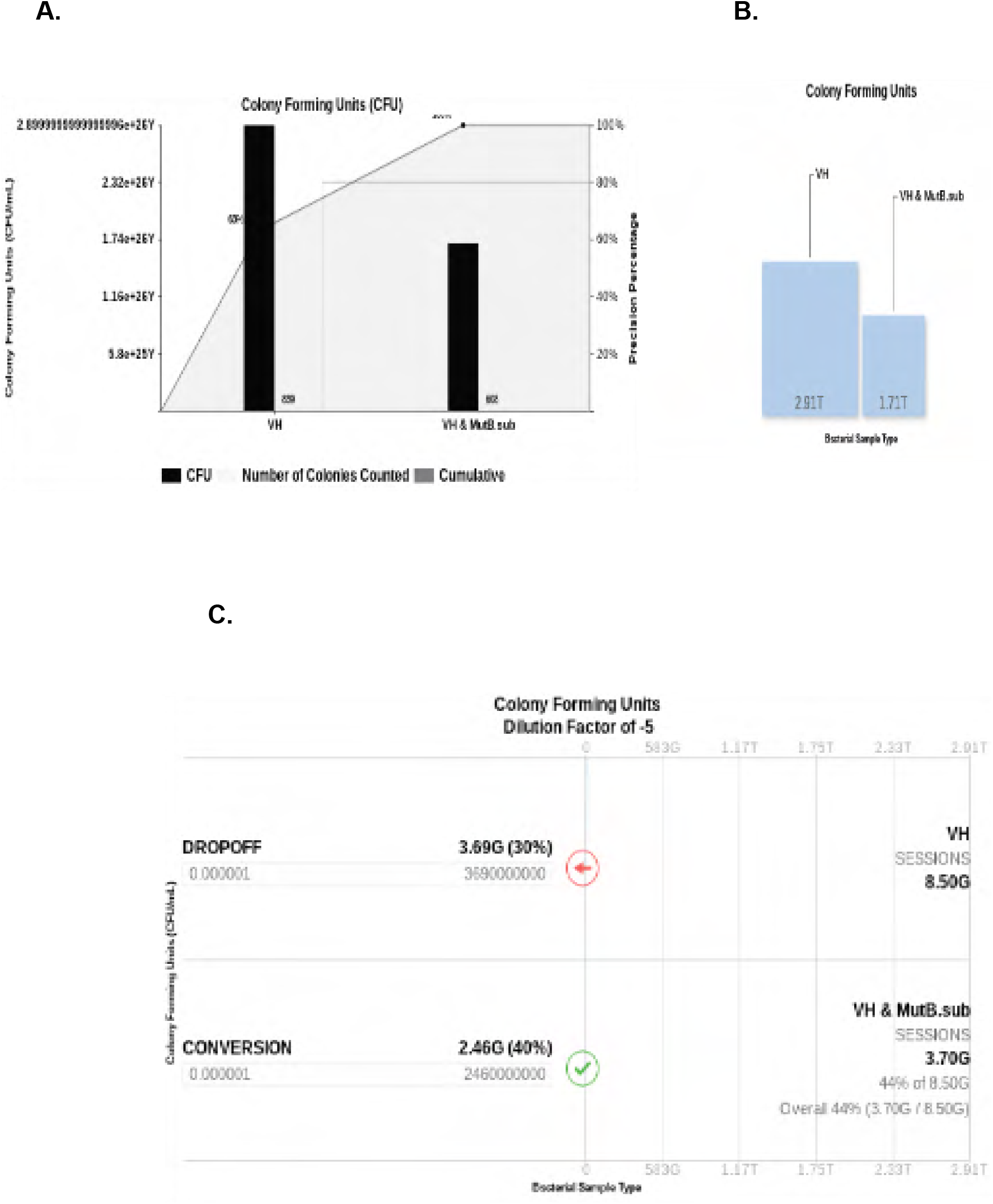
Colony Forming Units (CFU) Analysis. **A.** V.H. had a total CFU of 2.89E26 as Δ*B.subtilis* mutant-V.H co-cultures produced lesser CFU values. **B.** V.H outranked and overexpressed bacterial growth at 291T of CFU compared against Δ*B.subtilis* mutant-V.H with a 171T CFU. **C.** V.H had 8.50G CFU values as Δ*B.subtilis* mutant-V.H yielded a 3.70G CFU after a dropoff and conversion of 0.0000001.

### Biofilm formation was amplified in Δ*B.subtilis* mutant-VH co-cultures, but reduced in *E. Coli* co-cultures

Biofilm is vital for the survival, persistence, and for the proliferation of bacterial cells. Biofilm serves to insulate and protect bacterial cells from external antimicrobials such as antibiotics and other antagonistic microbes. Biofilm is also a major contributor to the increased virulence and proliferation of pathogenic bacteria, and it also invigorates antibiotic resistance. For this reason, the effects of Δ*B.subtilis* mutants on biofilm formation were measured and analyzed via biofilm assays of Δ*B.subtilis* mutant *V. harveyi and Ecoli* co-cultures. The biofilm production of *V. harveyi* and *E. Coli* co-cultures with wild type *B. subtilis* were compared to the Δ*B.subtilis* mutant *V. harveyi and E Coli* co-cultures. The wild type bacterial strains of *V. harveyi* and *E. Coli* were used as control samples for comparative analysis. For *V. harveyi* samples grown with Δ*B.subtilis* mutants, biofilm formation exceeded the levels of biofilm generated by *V. harveyi* combined with WT *B. subtilis*.

A Likert scale ranging from 1 to 5 was used to quantify the intensity of crystal violet absorbance added to the samples. A score of 4-5 equaled a high intensity of crystal violet coloration, and a value of 3 to 4 represented a moderate intensity of crystal violet absorbance. If a sampling well did not appear with much coloration or no biofilm formed, these samples were given a score of 0 to 1. The Δ*B.subtilis* mutant-VH co-cultures averaged a score of 3.9, and the WT *B. subtilis-V.H* samples averaged a 3.3 crystal violet absorbance score. The control or the WT *V. Harveyi* averaged a 3.8 CV intensity score. WT *B. subtilis-V.H* samples, the Δ*B.subtilis* mutant-VH co-cultures, and the controls each scored a total of 40, 47, and 46 of CV intensity, respectively (Figure 4). The p-Values, after performing a Two-Way ANOVA analysis of each group’s CV scores, computed to 0.134, 0, and 0.64 for comparing the difference between the scores, the difference between the bacterial sampling groups, and for determining the variable difference between the bacterial groups and their CV scores. There was an increase in biofilm yielded when culturing The Δ*B.subtilis* mutant-VH treatment groups versus the WT *B. subtilis-V.H* sample groups. However, the CV scores, for the Δ*B.subtilis* mutant-VH co-cultures, of biofilm formed were equally close to the WT-V.H sample scores. There seemed to be evidence of a cooperative or competitive interaction between the Δ*B.subtilis* mutant and WT *V. harveyi* during co-culturing, which resulted in amplified biofilm formation. Biofilm formation decreased after co-culturing *E. Coli* wth Δ*B.subtilis-*CRISPR-accA-mutants with a p-Value of 0.0001 (Figure 5). Therefore, to further analyze this possible competitive or cooperative synergy between Δ*B.subtilis-*CRISPR-accA-mutants with *V. harveyi* and *E. Coli*, Δ*B.subtilis*-mutant-V.H antimicrobial activity was determined via the cross-streak method.

**FIGURE 4.**
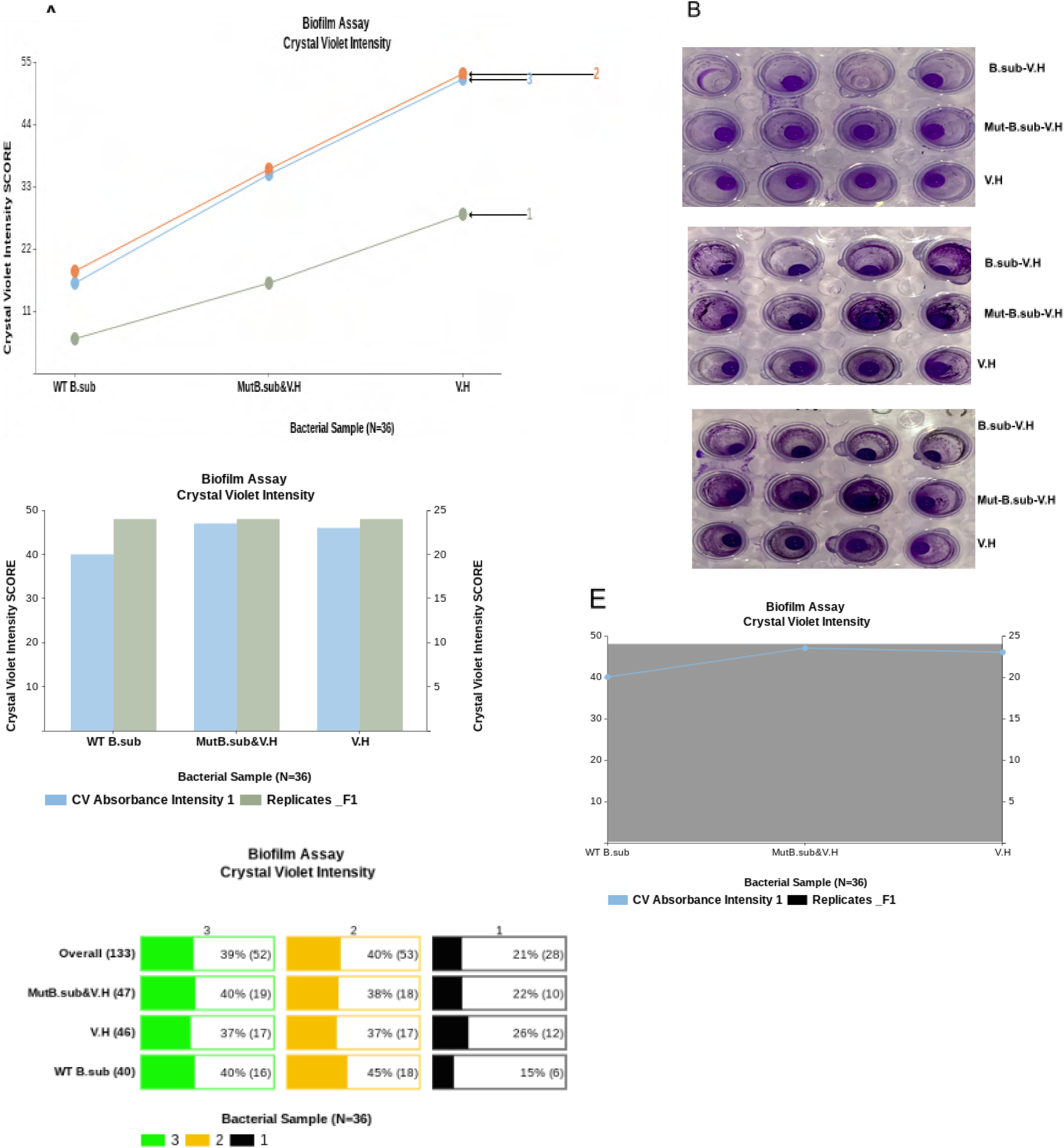
Biofilm Assay. **A.** Δ*B.subtilis* mutant-V.H and V.H showed a higher score of CV intensity than the WT-*B. subtilis-V*. H co-cultures. **B.** The crystal violet stain of Δ*B.subtilis* mutant-V.H appeared with the highest intensity. **C.** Δ*B.subtilis* mutant-V.H co-cultures showed a higher intensity of Crystal Violet absorbance with a total score of 47. **D.** The Biofilm assays were performed in triplicates with replicate 2 exhibiting the highest percentage of scored biofilm formation at 40%. **E.** Δ*B.subtilis* mutant-V.H had the highest expression of CV absorbance intensity.

**FIGURE 5.**
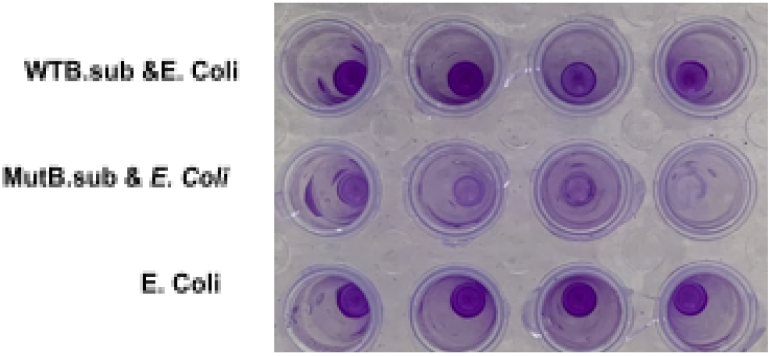
Biofilm Assay for *E. Coli* and Δ*B.subtilis*-CRISPR-accA-mutants. E. Coli had a total score of 16 with an average CV intensity of 4 (Avg=4, SD=0). Δ*B.subtilis*-CRISPR-accA-mutants combined with *E. Coli* averaged a score of 1 CV intensity and a total score of 4 (Avg.=1 SD=0.8). *E. Coli* grown with WTB.*subtilis* averaged a 3.5 CV score and had a total score of 14 (Avg=3.5, SD=0.58).

### The Cross-Streak of CRISPR-accA-Mutant *B. subtilis* with *V. harveyi* and *E. Coli* lacked antimicrobial activity

A single swab of Δ*B.subtilis* mutant LB liquid cultures in the center of the agar plates were cross streaked with WT *V. Harveyi* and *E. Coli*. The cross-streak was used because it is an efficient method for detecting antimicrobial effects of a bacteria on a specific target bacterial strain. The results from the cross-streak showed no antimicrobial activity on the WT *B. subtilis-V.H* plates or on the Δ*B.subtilis* mutant-V.H plates. There were no zones of inhibition to be quantified. There were no present zones of inhibition between *V. harveyi* or *E. Coli* when cross-streaked perpendicular to Δ*B.subtilis* mutants (Figure 6). Next, to detect any additional enhancement of antimicrobial activity, the levels of antibiotic resistance in Δ*B.subtilis* mutant-V.H co-cultures were tested by Kirby-Bauer Disk Diffusion tests.

**FIGURE 6.**
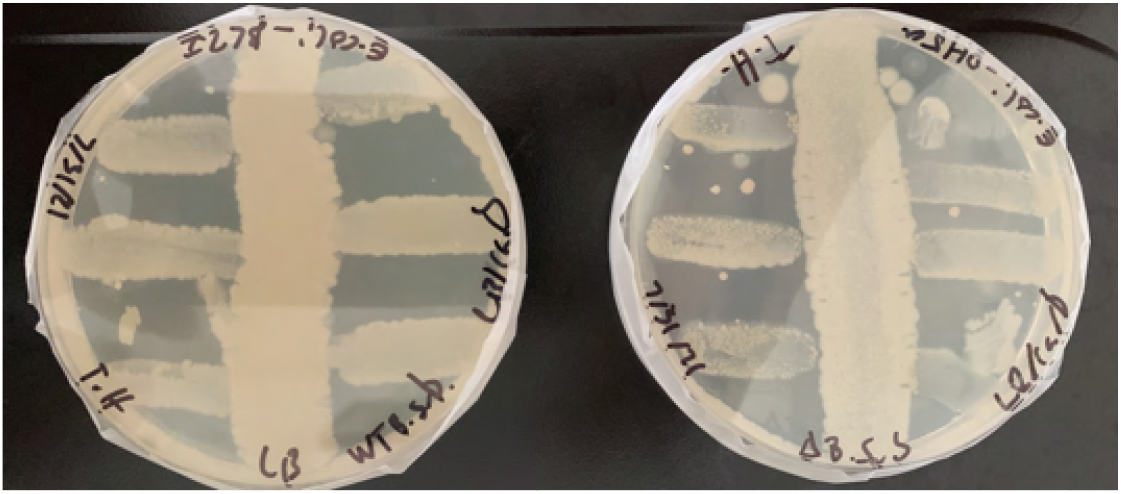
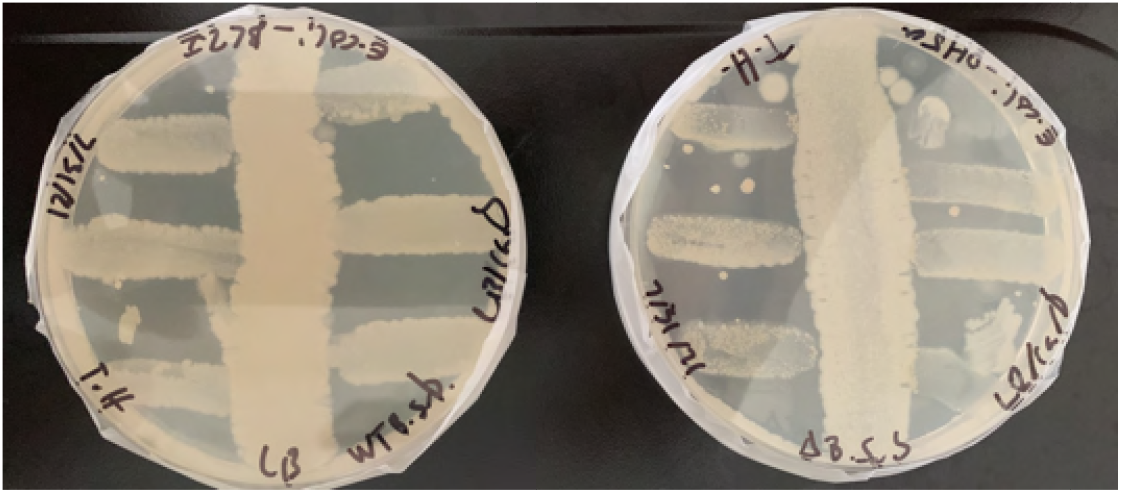
Cross-Streak Method and Antimicrobial Activity. The top plates include the Δ*B.subtilis* mutant with V.H and WT B. subtilis with VH cross streaks. The bottom plates are the cross-streak results from Δ*B.subtilis* mutants and WT B. subtilis with E. Coli. No antimicrobial activity was exhibited from the cross-streaked platΔ***B.subtilis* mutant-V.H and Δ*B.subtilis* mutant-*E. Coli* co-cultures showed more antibiotic resistance than the control samples**

Antibiotic resistance was measured in order to examine the antimicrobial potential of CRISPR-Cas9-accA-Δ*B.subtilis* mutants. The results of the antibiotic resistance further complicated the expected observation that Δ*B.subtilis* mutants would substantially decrease the antibiotic resistance of *V*. and *E. Coli*. The antibiotic resistance of Δ*B.subtilis* mutant-V.H and WT *V. harveyi* cultures were closely identical (Figure 7). In fact, the Δ*B.subtilis* mutant-V.H cultures seemed to share the exact amount of antibiotic resistance with nearly equal zones of inhibition quantities (Figure 8A). The Δ*B.subtilis* mutant-V.H, the WT-V.H, and the control samples were all sensitive to chloramphenicol and to kanamycin. All co-cultures were resistant to carbenicillin and moderately resistant to tetracycline (Figure 8B). The antibiotic resistance of Δ*B.subtilis* mutant-V.H co-cultures compared against the control samples had a Two-Tailed p-Value of 0.51 after using a paired t-test. The *E. Coli* samples showed similar levels of antibiotic resistance as the Δ*B.subtilis-mutant-E. Coli* co-cultures. All Δ*B.subtilis* mutant-*E. Coli* sample groups were more resistant to each antibiotic. The p-Values equaled 0.49 for Δ*B.subtilis* mutant-*E. Coli* compared against *E. Coli* after a One-Way ANOVA analysis. The antibiotic resistance levels of the co-cultures seemed to resemble and were identical to the antibiotic resistance expressed by the control sample groups, which included *V. harveyi* and *E. Coli* (Figure 9).

**FIGURE 7.**
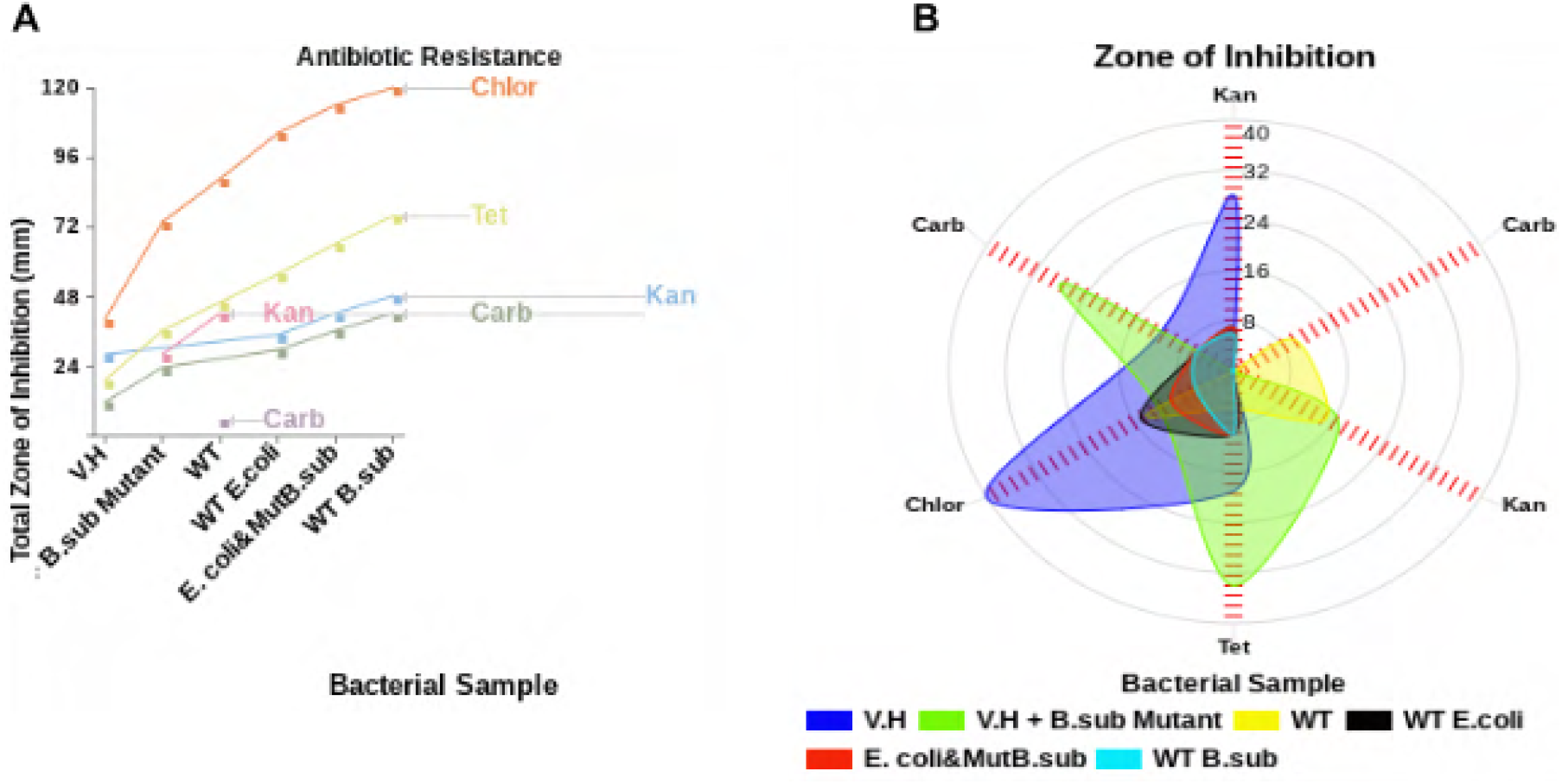
The Antibiotics of the Co-cultures. **A.** All the samples were resistant to carbenicillin. Group samples were more sensitive to chloramphenicol. Δ*B.subtilis* mutant-*E. Coli* was more sensitive to tetracycline while Δ*B.subtilis* mutant-V.H was moderately sensitive to kanamycin. **B.** Δ*B.subtilis* mutant-V.H was more resistant to chloramphenicol than WT *V. harveyi. ΔB.subtilis* mutant-*E. Coli* showed more resistance to chloramphenicol than WT E. Coli.

**FIGURE 8.**
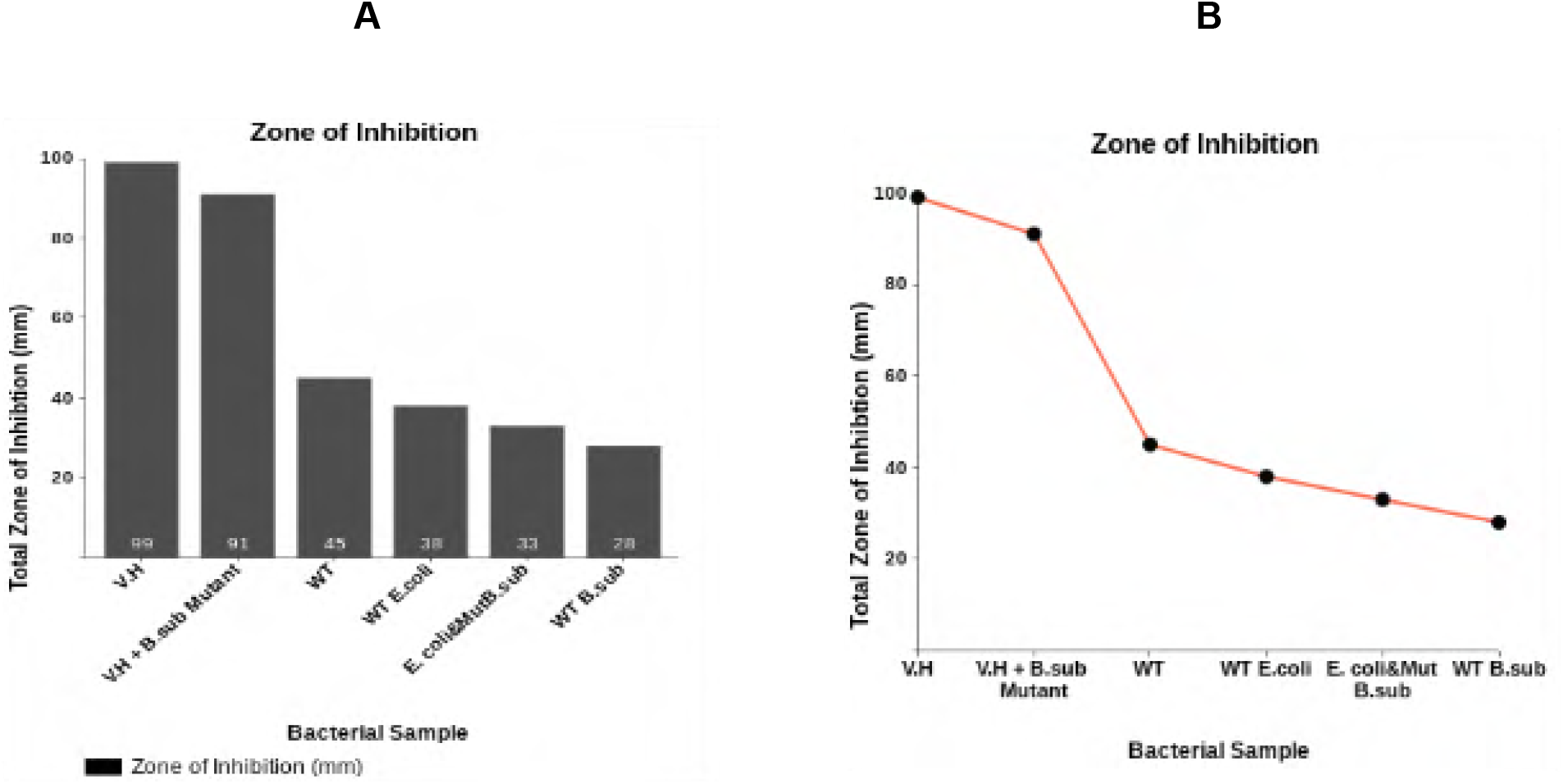
Zones of Inhibition (mm) The zones of inhibition were measured in millimeters. **A.** The zone of inhibition for *V. harveyi* totaled to 99 and 91 for Δ*B.subtilis* mutant-V.H. WT *E. Coli* had a total zone of inhibition of 38 as Δ*B.subtilis* mutant-*E. Coli* yielded a total ZOI of 33. **B.** Wild type *V. harveyi* and *E. Coli* samples showed increased antibiotic sensitivity, however, the Δ*B.subtilis* mutant-V.H and Δ*B.subtilis* mutant-*E. Coli* co-cultures slightly decreased in antibiotic resistance.

**FIGURE 9.**
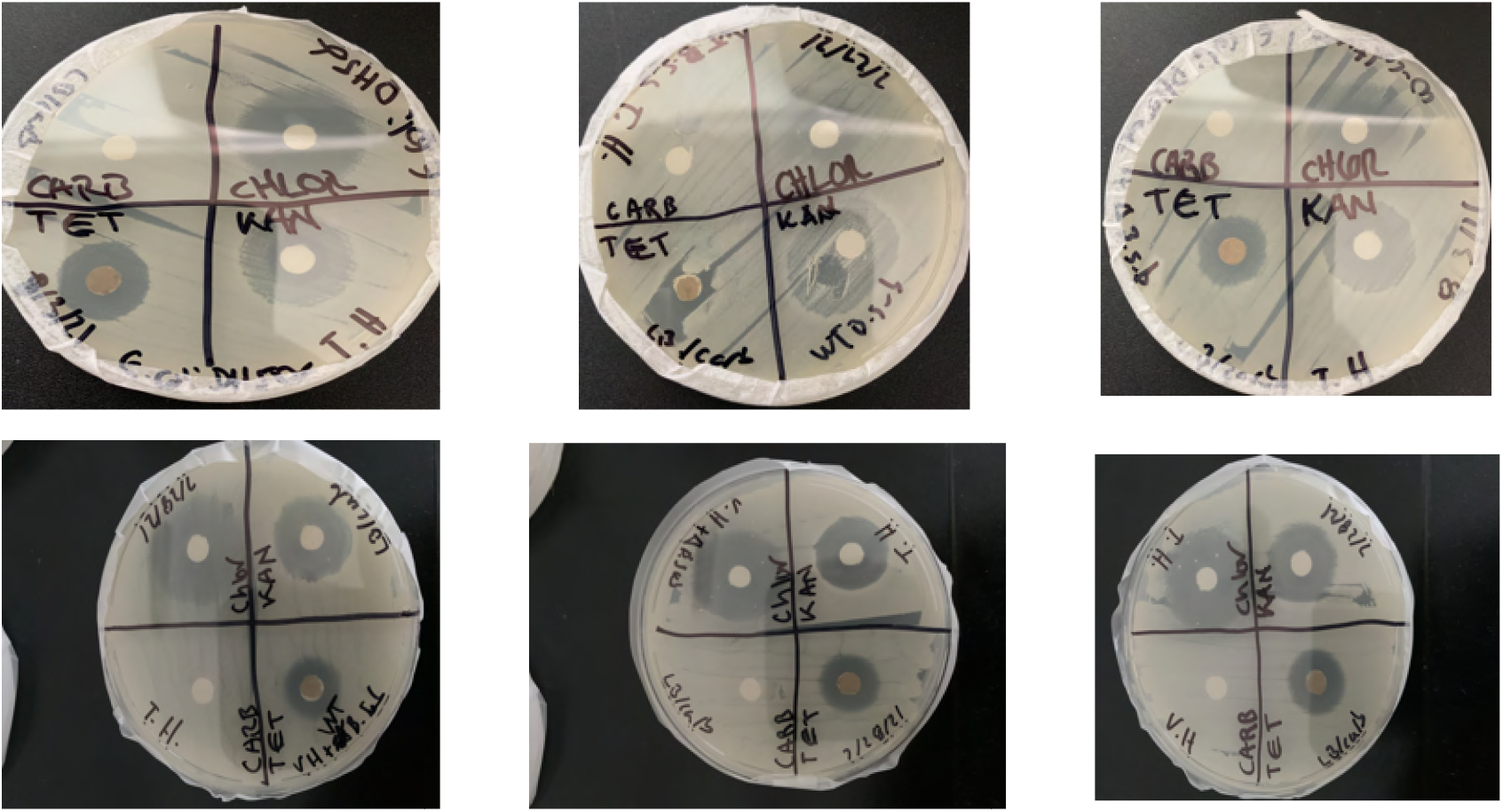
Antibiotic Resistance Tests. The top plates include the antibiotic resistant test results for *E. Coli* co-cultures and the bottom plates represent the results for *V. harveyi* sample groups. The top plates showed more moderate sensitivity to resistance of the antibiotics carbenicillin, Chloramphenicol, Tetracycline, and Kanamycin. The WT *E. Coli* was more sensitive to tetracycline, chloramphenicol, and to kanamycin than the WT B. subtilis and the Δ*B.subtilis* mutant-*E. Coli* co-culture. *V. harveyi* on the bottom plates showed identical and similar antibiotic resistance and sensitivity as the Δ*B.subtilis* mutant-V.H and WT B. subtilis co-cultures. All three bottom plates were resistant to carbenicillin and tetracycline, moderately sensitive to kanamycin, and sensitive to chloramphenicol.

## DISCUSSION

Probiotics have become popular, however, the effectiveness of probiotics is still conflicted with different opinions in the industry, in medical, and in scientific communities [9]. A method of precision is needed to form probiotics that bridge this gap between probiotic strains, individuals, and their microbiome [9]. There is an enlarged source of literature that conflicts with evidence-based clinical protocols for using probiotics [9]. Probiotic strains interact with the gut microbiota by competing with pathogenic bacteria for nutrients to give support to the homeostasis of microbiota [10]. Probiotic strains dispel other microbes through antagonism. The antagonism of probiotic strains involves their production of organic acids and of bacteriocins [10]. The organic acids and bacteriocins can actively combat pathogens in the human urinary tract and in human and animal gut. For example, Bifidobacteria produce acetate and can feed other bacteria in the gut microbiome.

Probiotics can prevent and inhibit the growth of pathogenic bacteria, amplifying human health [11]. Probiotics provide protection by competing for nutrients necessary for growth and for proliferating at higher rates than pathogens. Many studies have proven that *Lactobacillus rhamnosus* strains, GG and *L. plantarum* can attach to enteropathogenic *E. Coli* while in the GI tract [11]. The interest in probiotic bacteria is increasing specifically for the creation of foods. In a study by Tarrah et al. 8 *Lactobacillus* strains were isolated from infant feces and were tested for signals of probiotic properties such as antimicrobial susceptibility, hemolytic activity, and any inhibition of biofilm formation in other bacteria [12].

Increasing the concentration of probiotics led to increased microbial diversity in the GIT of shrimp [13]. The amplified bacterial diversity included increased levels of the *gemmatimonadetes, acidobacteria, deltaproteobacteria* and *xanthomonadales*. Mixing the species of probiotics promotes growth, optimizes immunity, and influences the microbiota in white shrimps [13]. The rate and kinetics of *V. cholerae* growth was inhibited in media enhanced with antibiotics and 7 different strains of *lactobacilli* [14]. In this study, genetically modified *B. subtilis*, a potential probiotic, was cultured with target bacterial strains, which affected bacterial growth, biofilm formation, antibiotic resistance, and antimicrobial activity of the co-cultures.

### Bacterial Growth

There is an enlarged interest in the probiotics of Bacillus species and in *B. subtilis* [5]. Bacillus species can prevent respiratory infections and gastrointestinal disorders such as irritable bowel syndrome. However, the pathway and mechanism in which Bacillus species function as probiotics is still unclear [5]. It seems that *B. subtilis* maintains an auspicious balance in the gut microbiota and improved the cell growth and potency of the LAB probiotic [5]. This study found that modifying *B. subtilis* to express CRISPR-Cas9 cleavage of gene *accA*, inhibiting fatty acid synthesis in target bacterial cells, lessened the bacterial growth of *V. harveyi*and the *E. Coli* strains. Co-culturing (-)*accA*-mutant *B. subtilis* with *V. harveyi* and *E. Coli* significantly reduced the CFU levels of the co-cultures because Gram-negative bacteria, such as *V. harveyi* and *E. Coli* are innately resistant to LAB bacteriocins because they have an outer membrane that is a physical obstacle for bacteriocin uptake [15]. However, if the outer membrane can become destabilized, then Gram-negative bacteria can become sensitive to bacteriocins [15]. Lactic acid formulated and released by LAB can permeabilize the Gram-negative bacterial outer membrane and allow the influx of antimicrobials whereas more molecules can cross the bacterial membranes of Gram-negative bacteria [15]. The lowered CFU levels in the mutant *B. subtilis, V. harveyi* and *E. Coli* cultures may be attributed to the mutant *B. subtilis* sharing of plasmid CRISPR-Cas9-*accA* vectors that reduced FAS in the target bacteria, leading to the breakdown of the outer lipid membranes in the *V. harveyi* and *E. Coli*.

The mutant *B. subtilis* may have permeabilized the *E. Coli* and *V. harveyi* outer membranes, which increased the influx of *B. subtilis* antimicrobials further decreasing the CFU values of the mutant-to-target bacterial co-cultures. *B. subtilis* cells generate a vast diversity of substances that induce antimicrobial activity. Many *B. subtilis* cells produce lipopeptides such as surfactants, iturins, and penguins. These antimicrobials can operate as antibacterials, anti-virals, antifungals, and cause anti-tumor activities. They can disrupt bacterial structures by lessening the surface tension of biofilms and via quorum sense inhibition. The inhibited quorum sense also inhibits formation of biofilm [5]. Lipopeptides can signal the induction of biofilm formation in *B. subtilis* [5]. *B. subtilis* antimicrobials combined with permeabilization of the target bacterial outer membranes significantly decreased the CFU results of the co-culture assays.

### Biofilm

Biofilm is an organized community of microbials sheathed in a polymeric matrix as it is attached to an biotic or abiotic surface. Biofilms normally contain a mixed and diverse microbial community or communities. These microbial communities interact via communication mediated by signaling molecules. These interactions between interspecies cause mutualistic activities or effects of antagonism [16]. The normal microbiota protects against upper respiratory tract infections (URTIs). However, dysbiosis or an imbalance in microflora can cause increased colonization and infection by pathogenic opportunistic microbes [16]. The rate of dysbiosis depends on the ratio between the number of disease-associated pathogenic species and health promoting species [17].

Probiotics that produce outstanding levels of biofilm can greatly inhibit the colonization of pathogenic bacteria in the gut epithelium. Likewise, the biofilm production, in this study, was amplified in mutant *B. subtilis* and *V. harveyi* co-culture because probiotic bacteria can generate biofilms, and they can efficiently adhere to colonic cells [14]. Every probiotic isolate in the Kaur et al. study could strongly adhere to colonic cell lines such as HCT-15 [14]. Through the auto-aggregation properties of probiotic bacteria, bacterial micro-colonies can be formed, biofilms can mature, and this process can release exopolysaccharides that cause biofilm maturation. Every lactobacilli showed more than 90% of auto-aggregation after eight hours [14]. After bacteria attach, their autoaggregation and swarming motility accelerate and more planktonic cells present increase colony formation [18]. These activities, during the biofilm generation process, induce the maturation of biofilm [18]. Because probiotics can form biofilm, probiotics can initiate effective colonization and successfully treat dysbiosis, this fortifies probiotics to survive many different environmental conditions. Probiotics, as a consequence, can cause colonization and increased survivability of its population and species [18]. Probiotics were found by Schwendicke et al to decrease growth and biofilm formation of cariogenic-like bacteria, which includes *Streptococcus mutans* [19].

Biofilms protect against pathogenic bacteria that can avoid host defenses via a protective shell. Biofilms are a favored habitation for pathogenic bacteria to produce virulence whereas biofilms of healthy tissues can signal the presence of a damaged gut [20]. That is, for example, *E. Coli* cells can form biofilms, which develop into ulcerative colitis. However, if biofilms are produced in a healthy gut, functions of microbiota can benefit host health by improving host defense responses [20]. When comparing the mutant *B. subtilis* cultured with *E. Coli* against the wild type *B. subtilis* with E. Coli, the CRISPR-Cas9-accA modified *B . subtilis with E. Coli* produced a stronger antimicrobial response against *E. Coli* than the wild type *B. subtilis*. These significant results demonstrate that the antimicrobial activity of CRISPR-Cas9-accA modified *B. subtilis* was enhanced partly because of the biofilm production of the mutant *B. subtilis-E. Coli* mixed culture significantly decreased more than the wild type *B. subtilis-E. Coli* mixture.

A possible hypothesis for this observation, of mutant CRISPR-Cas-*accA B. subtilis* having an enhanced antimicrobial response against *E. Coli*, is that deleting accA permanently inactivated the fatty acid synthesis in *E. Coli*, and after the stationary phase, where most metabolic activity rates are lower during the stationary phase, *E. Coli* could not re-initiate FAS, [21]. *E. Coli* during the stationary phase has a time to rest and rebuild by storing and accumulating pyruvate that were exhausted during the amplified metabolic activity of the growth phase. However, when sufficient amounts of carboxyltransferase (CT), consisting of AccDA, are available, the enzyme remains bound to the transcript and further inhibits the synthesis of proteins like a negative feedback loop [22], but after the stationary phase, the availability of CT may be consumed and FAS in *E. Coli*, without accA gene expression, may have remained stagnant in a negative feedback loop, not producing sufficient levels of fatty acids to restore the cell membrane for early biofilm formation, thereby heavily reducing the biofilm.

The biofilms in a healthy gut optimize the sharing of nutrients between microflora and their host, increasing the survival of gut bacteria [20]. Probiotic biofilms greatly benefit and enrich the health of the host through increased colonization and long residence in the gut mucosa, and this lessens pathogenic bacterial colonization. Antibiotics combined with seven different isolates of *Lactobacillus* caused more than 90% of decreased *V. cholerae* biofilm formation [14]. Because lactobacilli can form many fortified biofilms, lactobacilli can survive and proliferate *in vivo*. When biofilms are created, they become resistant to antibiotics where many antibiotics only combat planktonic *V. cholera* cells and cannot limit biofilm-dispersive activity [14]. As a result, many probiotic strains possess antimicrobial and antibiofilm capabilities against *V. cholerae* and can be effectively and clinically administered. A current model system was used to show how *B. subtilis* enhances biofilm growth via its shared growth with LAB [5]. The model system demonstrated the increased protection of LAB from increased heat and from the acidic regions of the gastrointestinal tract, in which the LAB probiotics traversed when added to *B. subtilis* [5].

### Antibiotic Resistance

*Bacillus* strains reside in soil, air, in fermented foods, and in the gut of humans. Probiotics of *Bacillus* produce spores that germinate and grow into sporulation in the GI tract. These spores allow the survival of *Bacillus* bacteria to persist [23]. However, Bacillus strains can transfer and share antibiotic resistance genes, produce enterotoxins, and biogenic amines. Probiotics of *Bacillus* can become symbiotic organisms in the host, temporarily [23]. However, strains of *Bacillus* are resistant or susceptible to all antibiotics, secluding and excepting ampicillin [23]. As a result, this study showed increased antibiotic resistance in mutant *B. subtilis* with *V. harveyi* and *E. Coli* co-cultures, which is indicative of the transfer and sharing of antibiotic resistance genes between *B. subtilis* and the targeted bacterial strains.

Assessing the strains of next generation probiotics (NGPs) is required, needed, and highly pertinent [24]. The issue for the safety of NGPs includes their ability to carry, share, and spread antibiotic resistance genes [24]. In addition, NGP species are extremely difficult and arduous to produce and manufacture where many NGPs will only be purchased as supplements [24]. The safety of many new and novel probiotics offered to consumers requires cautiously choosing less toxic probiotics that are most beneficial to human health. Probiotic bacteria can vertically transfer genes with other commensals and pathogens in the lower gastrointestinal tract and this is a major cause of concern for probiotic safety [25]. Combining probiotics with antibiotics when administering antibiotic therapy has been proven to restore health [Selvin]. However, horizontal gene transfer of multidrug resistance unto other pathogenic bacteria and to commensals presents a major threat to the microflora of the lower intestines [Selvin]. Using probiotics with antibiotics simultaneously causes many resistant activities to develop that override the antimicrobial effects of the antibiotics applied [25].

Probiotics will share in transferring antibiotic resistant genes between pathogenic and commensal bacteria in the lower GIT [25]. The likelihood of this exchange of drug-resistant genes amplifies our need to further research the safety of the bacterial strains inside probiotic supplements [25]. Das et al. agrees that drug-resistance genes transferred between probiotics and other bacteria is a source of enormous concern but a huge lack of information is still not currently provided [26]. Many research studies and reports have supported the design and formulation of commensal bacteria into next-generation probiotics, but admit that this should be more cautiously pursued [27]. Probiotics can restore balance in the microflora and in the microbiota of the intestines after antibiotic therapy, can colonize the lower GIT, and eradicate pathogens. However, antibiotic resistant genes can accumulate in the gut, thereby eliminating many of the beneficial properties of probiotics [27]. Because the transfer of resistant genes frequently occurs in mouse models, it also must occur in building a reservoir of antibiotic resistant genes in the human gut [27]. Li et al. recommends and provides an approach for further research in developing and applying probiotics [28]. Lit et al. found that the potential risks of using probiotics include increased pathogenicity, infectivity and over production of immune responses [28]. Probiotics enlarge the resistome in the GI mucosa through mediating the spread of strains harboring vancomycin resistant genes but without the transfer of resistant genes expressed in probiotic strains [29]. Stool samples in the research study by Montassier et al. showed through direct sampling that combining use of probiotics with antibiotics affects the gut resistome [29]. Current published clinical trials showed the significance of implementing patient-individual-specific profiling of metagenomics of the GIT resistome via direct sampling [29].

### Antimicrobial Activity

Probiotics such as Lactobacillus inhibited enteropathogenic growth, showing inhibition zones between 12 and 32 millimeters [30]. The four Lactobacillus strains in the Tebyanian et al. study presented many potential antimicrobial molecules to antagonize enteric pathogens in humans [30]. They recommended further studies of the mechanisms that allow probiotics to improve human health [30]. A study by Lashani et al. found 15 isolates were inhibiting the growth of foodborne pathogenic bacteria while 24 other isolates lacked any inhibitory signaling effects [31]. Five probiotic strains inhibited the growth of *E. Coli* pathotypes where all chosen strains had a great antimicrobial effect against *E. Coli* O157:H7 strains, but did not inhibit the growth of Enterohemohagic *E. Coli* [32]. Karimi et al. suggested and encouraged the use of probiotics to treat disease [32]. However, the present study did not detect any antimicrobial activity of the mutant *B. subtilis* with the target bacterial co-cultures, using the cross-streak methods. No antimicrobial activity was detected because the *V. harveyi* and *E. Coli* bacterial strains co-cultured with the mutant *B. subtilis*, were non-pathogenic. Probiotics may have a greater propensity to secrete antimicrobial compounds when pathogenic microbes are present. Therefore, the findings of antimicrobial activity via the cross-streak method, in this study, was limited due to lack of access to pathogenic target bacterial strains. Nevertheless, further research may be needed to elucidate the interaction between probiotics, non-pathogenic bacteria, and with other commensals.

Many LAB strains have indirectly mediated adverse effects when used to eliminate pathogenic microbes in a host and also via LABs’ antagonism of unfavorable microorganisms [33]. The first defense against bacterial infections in the GIT is antibiotic therapies. When the microbiota in the gut is unbalanced, as a result of antibiotic use, the microbiota can remain unbalanced [34]. Narrow-spectrum antibiotics, changes to diet, transplanting fecal matter, and use of probiotics can restore balance and diversity to the microbiota of the intestines and enhance the health of the host [34]. Probiotic species such as *Bifidobacterium* and *Lactobacillus* have been recommended to improve the health of patients struggling with intestinal infections. However, the mechanisms of probiotic properties are not completely understood [34].

Carzorla et al. support the uses of L. casei CRL 431 and L. paracasei CNCM I-1518 because these probiotics are safe, potent, and frugal for amplifying antimicrobial activity in the intestines [35]. Lactic acid bacterial species can release their lactic acid in a dose-like approach, eliminate pathogens, lower the attachment and colonization of pathogens, deactivate toxins while competing with the pathogens for a lack of present nutrients [35]. Bohora et al. demonstrated that a paradigm shift was uncovered in their study, in which the emphasis for total eradication of pathogens was changed into restoring the balance of the microbiome and the microbial environment within a host [36]. The probiotic effect depends on the bacterial strain; therefore, the genotypes and phenotypes of probiotics should be determined where these probiotics are safe, non-toxic, and cannot lead to pathogenesis [37]. Probiotics should not have factors of virulence and should lack acquired AR genes.

## CONCLUSION

Global probiotics were valued at 48.88 billion USD in 2019 and is projected to increase to 94.48 billion USD by 2027, which is a compound annual growth rate (CAGR) of 7.9% [38]. There is a great need for increased research that clarifies which probiotic is the most effective for yielding the best health results and to gather evidence from the people with the greatest response to specific probiotic strains and doses [10]. Probiotics, prebiotics, and synbiotics used as supplements yield many promising data results to combat enteric pathogens. Probiotics can compete with pathogenic microbes, isolate pathogens, stimulate, and regulate the immune response of a host by inducing gene expression internal and external of the host gastrointestinal tract.

However, probiotics are normally delivered as dried cultures where the drying process can degrade the cell’s structure, reduce viability, and lessen its potency [5]. During the drying process, an enormous amount of fluid is removed from the cells. This negatively affects the cells structure and physiology, resulting in cell death [5]. Another issue includes that most probiotic cells die during their shelf life, and also have difficulty surviving the transition through the GIT that is filled with stomach acid, degrading enzymes, and with bile salts of the intestines [5]. The nano-encapsulation of probiotic supplements are high cost and less accessible [39]. Therefore, a facile, frugal, accessible, and intrinsic method are needed to increase the health benefits of probiotic supplements [39].

## Acknowledgements

Special thanks are given to my mentors. They were always available to answer questions and provide support.

## Author Contribution

The author confirms sole responsibility for the following: study conception and design, data collection, analysis and interpretation of results, and manuscript preparation.

## Ethical Statement

This article does not contain any studies involving animals performed by any of the authors. This article does not contain any studies involving human participants performed by any of the authors.

